# The high abortion cost of human reproduction

**DOI:** 10.1101/372193

**Authors:** William R. Rice

## Abstract

Information from many large data bases and published studies was integrated to estimate the age-specific spontaneous abortion rate in an economically-developed human population. Accuracy was tested with published data from a diverse array of studies. Spontaneous abortion was found to be: i) the predominant outcome of fertilization and ii) a natural and inevitable part of human reproduction at all ages. The decision to reproduce is inextricably coupled with the production of spontaneous abortions with high probability, and the decision to have a large family leads to many spontaneous abortions with virtual certainty. The lifetime number of spontaneous abortions was estimated for a “canonical” woman (constrained to have average age at marriage, first birth, inter-birth intervals, and family size) in two populations: one with and the other without effective birth control (including free access to elective abortions). Birth control was found to reduce lifetime abortions more than 6-fold.

---

Abortion is one of the most socially contentious issues in the United States and many other parts of the world. This overarching social influence underscores the need to fully understand the intrinsic scope and role of abortion in human reproductive biology. To date there is only one statistically rigorous estimate of the age-specific rate of spontaneous abortion in humans (that includes both clinically detected and earlier, occult [unperceived] abortions), and this was based on women living in an economically-undeveloped country [1]. The sample size was large for a single study (all or part of 329 pregnancies were assayed): nonetheless this level of sampling represents an average of only 11 pregnancies screened per yearly age class (spanning 30 reproductive years). The purpose of this paper is to integrate recent data from many disparate sources to assess the rate of spontaneous abortion in a near-contemporary, economically developed population. With this synthesis I will: i) estimate the age-specific rate of spontaneous abortion in a European population with an exceptionally large (nation-wide) data base of medical records, ii) test the accuracy of this estimate with data from independent studies, iii) quantify the age-specific cost–in the currency of number of spontaneous abortions– of producing a newborn, iv) evaluate the influence of modern birth control–including elective abortions– on the expected number of abortions per lifetime.

## Results

The potential for spontaneous abortions to strongly influence the age-specific pattern of human female fertility is immediately apparent when comparing plots of age-specific fertility rate and the rate of clinically detected spontaneous abortions (e.g., [2]). In Fig. 1, I have plotted two reproductive metrics against age: i) the estimated yearly probability (as a percent) that a married woman delivers a newborn, and ii) the estimated probability (as a percent) that a clinically detected pregnancy spontaneously aborts. The fertility data are taken from a natural-fertility population in which modern contraception was virtually absent (frontier Mormons from 1850-1879, N = 17,806 women, reported in [3]), and the spontaneous abortion data are from Denmark between 1978-1992 (N = 1.2 × 10^6^ pregnancies, reported in [4]) – a population that has free modern birth control (including free-on-request elective abortions up to 12 weeks of fetal development) as well as a national registry of medical records for all citizens that records births and abortions. Ideally, the graph would include birth and abortion data from the same natural-fertility population, but these populations do not have the large data bases (that combine widespread medical testing and record-keeping) needed to accurately monitor age-specific spontaneous abortions. Despite the women coming from different continents, time periods and socioeconomic and nutritional conditions, and the difference in units (probability of a newborn per year vs. probability of an abortion per pregnancy) there is a virtually perfect negative correlation between births and abortions. This strong pattern has motivated the hypothesis that spontaneous abortion plays a predominant role in shaping the age-specific pattern of human fertility (i.e., the Wood-Boklage-Holman hypothesis: [1,5-9]). That it is an elevated abortion rate of conceptuses from older eggs of older women that causes fertility decline with advancing age in human females–rather than deterioration of other aspects of a woman’s reproductive anatomy and physiology– is supported by data from assisted reproductive technology clinics. The capacity to carry an embryo (from in vitro fertilization) to birth declines precipitously with age only when a woman uses her own eggs: not when she uses eggs from a young egg donor (Figure 2; see also [9b]).

**Figure 1.**
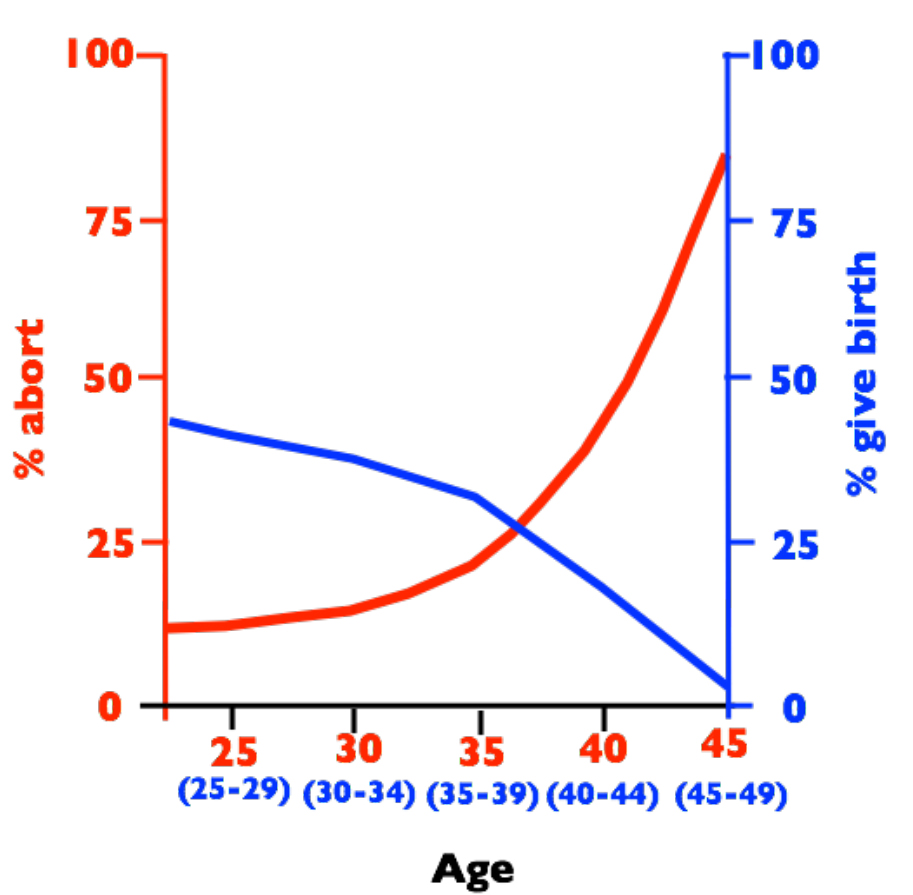
Birth rates per year (blue, percent women in each age class giving birth) and spontaneous abortion rates per clinically detected pregnancy (red, percent pregnancies aborting) vs. female age. Abortion rates are from a sample 1.2 × 10^6^ medical records from Denmark collected between 1978-1992 [4] and birth rates are from historical records of 17,806 frontier Mormons collected between 1850-1879 [3].

**Figure 2.**
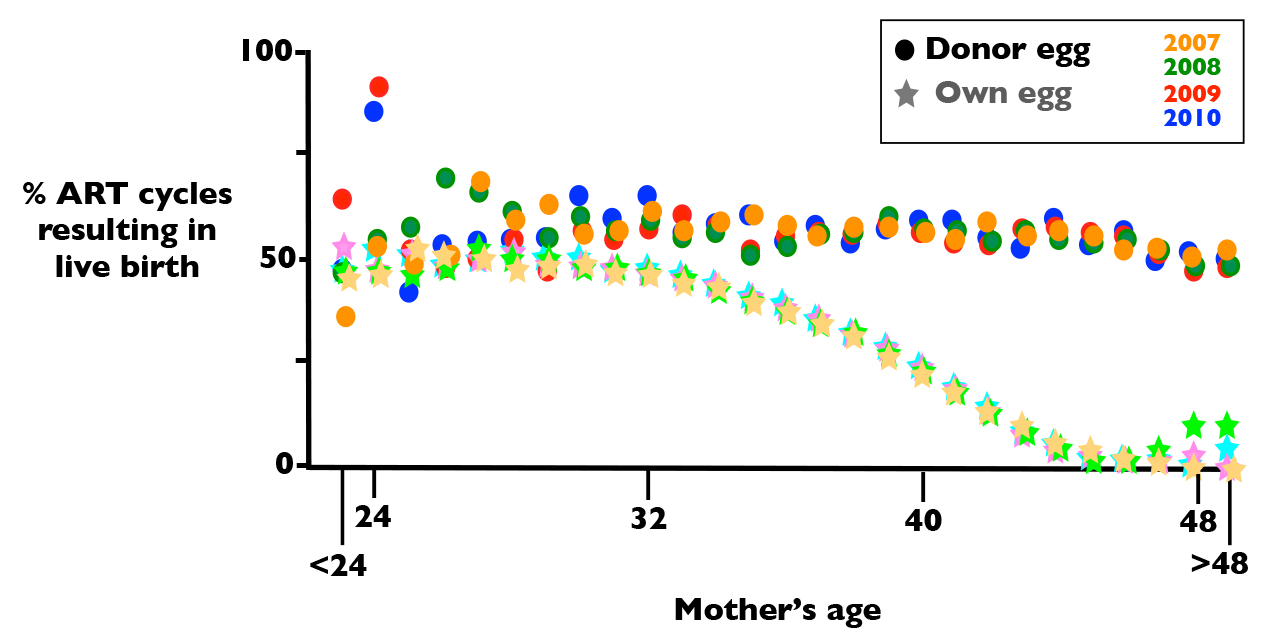
Age-specific success rate of pregnancies in women using assisted reproductive technology (ART) clinics using their own eggs (stars) or eggs from egg-donors (circles). Sample sizes: N_each year_ > 143,000 and N_total_ = 583,994. The age of egg donors is usually between 21 and 34 years old and the average age of egg donors in 2010 was 28 years old. Sample sizes for women less than 25 and more than 45 are highly rarified in the data set compared to ages between these extremes, and these data are therefore more variable. Data from the CDC web sitehttp://www.cdc.gov/art/reports/archive.html.

### Age-specific abortion rate

To explore a potential causative relationship between human female fertility and abortion rates, a complete measure of spontaneous abortion is needed, rather than the incomplete metric of clinically detected abortions shown in Fig. 1. Because I will focus here almost exclusively on spontaneous abortions (rather than elective abortions), henceforward I will use the term “abortion” to denote spontaneous abortion unless specified otherwise. The data on abortion shown in Fig. 1 forms an exceptionally strong empirical base for age-specific abortion rate because of the very large number of pregnancies (1.2 × 10^6^) that were monitored to construct it. Because of its uniquely large sample size, and hence low statistical sampling error, I will use this database as a foundation on which to construct a more complete measure of age-specific abortion rate in an economically developed human population.

The abortion data shown in Fig. 1 was based only on abortions in women who were hospitalized (and therefore the abortion was reported to a national registry of medical records). Andersen et al. [4] estimated that 20% of women experiencing a clinically detected spontaneous abortion in Denmark were not hospitalized during the period between 1978 and 1992, and a similar rate was reported during the same time period in a neighboring Scandinavian country (Finland, [10]). In Fig. 3, I adjust for missing abortions in Fig. 1 (due to lack of hospitalization) by adding one additional abortion for every four recorded abortions used to construct the curve in Fig. 1. The lowest curve (labeled “1”) is the unadjusted rate and the next higher curve (labeled “2”) is the hospitalization-adjusted rate.

**Figure 3.**
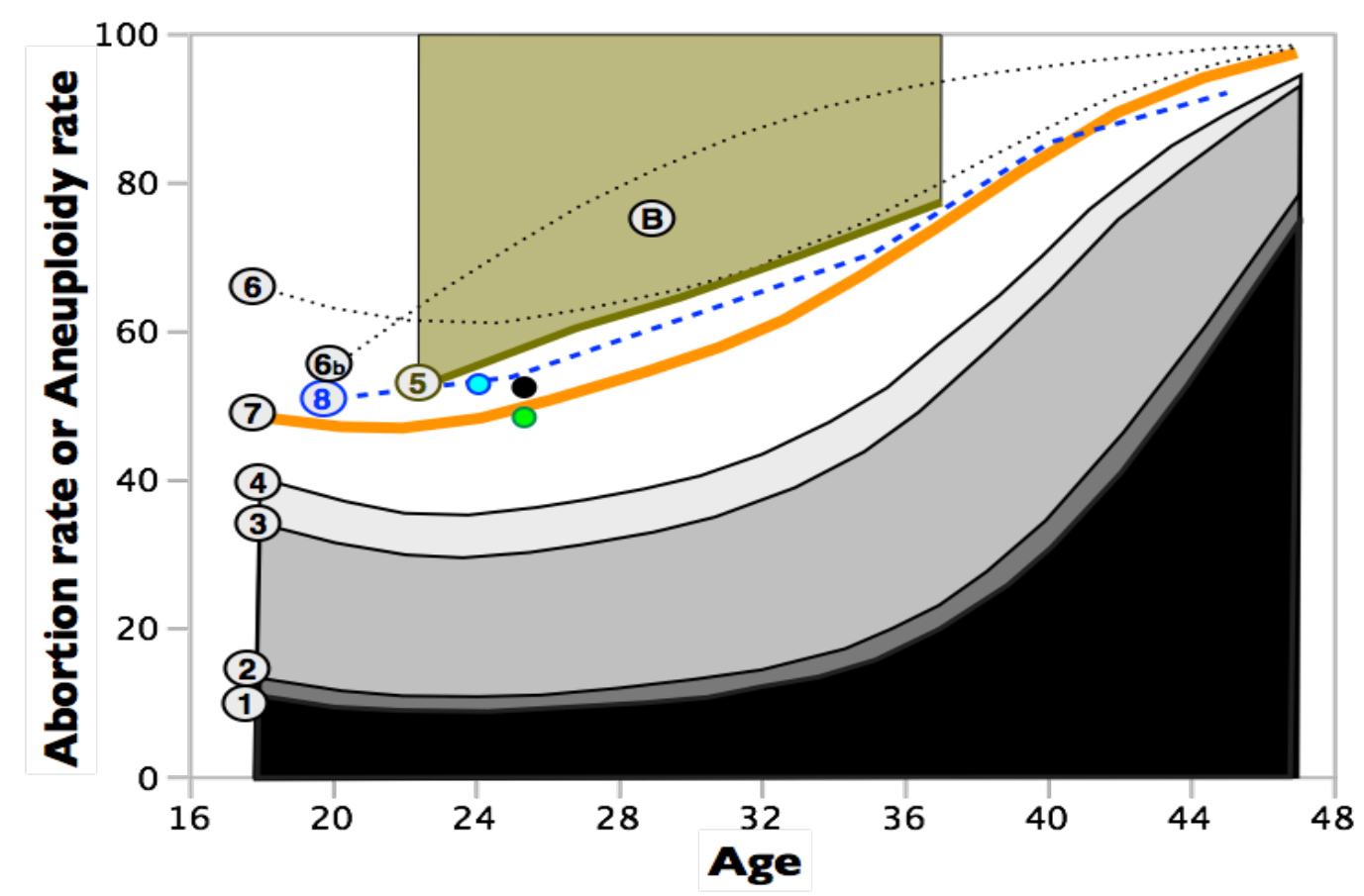
Estimated age specific abortion rates, with varying levels of downward or upward biases, and comparisons to age-specific rates of unbalanced karyotypes in early-stage embryos. All rates refer only to spontaneous abortions. Curve-1 is taken from Fig. 1 and represents clinically detected abortions in a large sample of women with clinically detected pregnancies and hospital records for abortions [4]. Curve-2 adjusts curve-1 upward by accounting for women who had an abortion after clinical detection of a pregnancy but were not hospitalized. Curve-3 adjusts curve-2 upward for many of the abortions that occur before clinical detection (occult abortions) but after implantation. Curve-4 adjusts curve-3 upward by including some of the preimplantation abortions that are caused by chromosomal monosomies. Curve-4 does not adjust for all abortions that are not clinically detected and therefore represents an estimated lower bound (floor) for the true abortion rate. Curve-5 is an estimated upper bound (ceiling) for the abortion rate based the rate at which fertile women experience a clinically detected pregnancy when sperm are delivered at the optimal time. This curve is a ceiling because it assumes that all failures to become pregnant are due to occult abortion. Point-B is the estimated abortion rate [7] (clinically detected + occult) based on pooling diverse information from all time-points of gestation (average age of women in these studies was about 29 years). Curve-6 estimates the total abortion rate (occult + clinically detected) by adjusting curve-2 for the ratio of occult to clinically detected pregnancies from the Boklage [7] study. Curve-6b is the same as curve-6 except that the age-specific ratios of occult to clinically detected abortions are the maximum likelihood estimates from an exceptionally statistically rigorous study of a natural fertility population in Bangladesh [1]. This curve represents the best available estimate of age-specific abortion rates in an economically developed population. Curve-7 assumes that nearly all spontaneous abortions are due to karyotypically unbalanced embryos. The green point is the estimated proportion of karyotypically unbalanced embryos from a sample of 46 fertilized eggs collected from young women (averaging 25.4 years old) during normal ovulation i.e., with no chemical stimulation to induce the ovulation of multiple follicles [11]. The black point is a sample from the same women (N = 303 eggs) but with moderate ovarian stimulation [11]. The blue point is the rate of karyotypically abnormal embryos from a sample of 305 fertilized eggs from 24 young egg donors (mean age = 24.0) who received standard levels of ovarian stimulation [12]. Curve-8 is the rate of karyotypically unbalanced embryos from a large sample (N = 5,918) of cleavage-stage embryos obtained from 726 women across 70 North American assisted reproduction clinics [13].

After accounting for spontaneous abortions that do not result in hospitalization, the detected rate of abortion would nonetheless only be a metaphorical tip of a much larger iceberg if most abortions occurred before a pregnancy could be clinically detected (this time point was ~3 weeks post-conception during the time period covered in Figs. 1 and 3; [7]). In the 1980s, a succession of laboratory studies using highly sensitive assays for human chorionic gonadotropin (hCG, produced by the human conceptus starting at implantation [14], and rapidly increasing above non-pregnancy levels as the implanted embryo develops) were used to detect pregnancies and abortions earlier than the standard clinical detection point (at ~2 weeks post conception; summarized in [1,7,8]). The “gold standard” study with the best design and appropriate controls was that of Wilcox et al. [15] in late 1980s which found that 31% of detected pregnancies aborted (62 abortions/198 pregnancies), with 9% after clinical pregnancy detection and 22% pre-clinical detection. To account for the preclinical abortions detected by studies like that of Wilcox et al. [15], I have added 22 occult abortions per 9 clinically detected abortions (adjusted for a 20% lack of hospitalization) to the graph in Fig. 3 (curve 3). Studies like that of Wilcox et al. [15] clearly demonstrate that most abortions are occult (occurring before a pregnancy is clinically detected) but could not measure preimplantation abortions nor those that occurred soon after implantation, so a substantial number of spontaneous abortions may have been undetected.

One way to further improve the estimate of the abortion rate is to account for at least some of the occult abortions that are not detected by high-sensitivity hCG studies, like that of Wilcox et al. [15]. A rationale for such an improvement comes from studies comparing aneuploidy in very young embryos (cleavage stage and blastocysts) to conceptuses that spontaneously aborted after a pregnancy had been clinically detected. At the early embryo stages, monosomy (missing one homolog from one or more of the 23 pairs of homologous chromosomes) and trisomies (a extra chromosome at one or more of the 23 pairs of homologous chromosomes) were found to be equally common [16-18]. But among conceptuses that survive to clinical detection, and then abort, essentially no autosomal monosomies were observed (<<1%; [19-22]). This repeated finding demonstrates that all autosomal monosomic conceptuses spontaneously abort before a pregnancy can be detected by elevated levels of hCG, i.e., prior to implantation. As a consequence, for every observed aborted conceptus that carried one or more trisomies after clinical detection, there must have been one occult aborted conceptus due to one or more monosomies. At least 50% of aborted fetuses are karyotypically abnormal and at least 60% of these carry one or more trisomies [22]. There are therefore at least 0.5^*^0.6^*^ = 0.3 occult abortions due to monosomy for every detected post-implantation abortion. In Fig. 3, I have added these additional occult abortions to the total (curve 4).

Curve 4 in Fig. 3 (with adjustments for non-hospitalization, some post-implantation occult abortions, and some pre-implantation occult abortions due to monosomies) represents an estimated lower-bound for the true abortion rate because it assumes that all implanted embryos that abort are detected and all mortality due to trisomy occurs post-implantation. To incorporate potentially undetected abortions and achieve a less downwardly biased estimate of total abortion, occult abortions must be estimated in a more inclusive manner. But before doing so, I will estimate an upper-bound for the age-specific abortion rate.

An estimated upper-bound for the age-specific abortion rate can be determined by measuring the maximum clinically recognized pregnancy rate per menstrual cycle for fertile couples with an optimal timing of sperm delivery. I focus on a large European study of couples that did not use effective birth control (no IUDs, diaphragms, condoms, nor birth control pills; [23]). There have been many studies estimating the maximum probability of a clinically detected pregnancy per menstrual cycle that includes sexual intercourse, but I focused on this study because it reports age-specific values, rather than a single value that was pooled across all ages. The sample size is substantial: 647 women, 2,939 menstrual cycles, and 433 detected pregnancies. The highest pregnancy rate occurred when couples had sexual intercourse two days prior to the estimated day of ovulation (estimated by both shifts in core body temperature and patterns of vaginal mucus discharge), and in this case the estimated probability of producing a detectable pregnancy was 0.53, 0.42, 0.36, and 0.29 for age categories (in years) of 19-26, 27-29, 30-34, and 35-39, respectively. The complementary probability for each of these values can be used to estimate the upper limit for the age-specific occult abortion rate. It is an upper limit because this estimate implicitly assumes that fertilization was 100% successful during each menstrual cycle that had an optimal timing of sperm delivery, so that all “no pregnancy” events were due to occult abortions. An upper bound for the total abortion rate is 1- (maximum pregnancy rate) + ([maximum pregnancy rate] ^*^ [proportion of the detected pregnancies that result a clinically detected abortion]), this latter term is calculated using curve 2 in Fig. 3. Using the mid-range of each age category, I have used these values to calculate the estimated upper bounds for total abortion rate (clinical + occult) in Fig. 3 (curve 5). The true (unbiased) curve for abortion rate vs. age is expected to be somewhere between the estimated upper ceiling (curve 5) and estimated lower floor (curve 4).

To obtain an unbiased estimate of the abortion rate, we need to add to curve 2 in Fig. 3 (probability of abortion given that a clinically detected pregnancy occurs) an unbiased estimate of total occult abortions. Boklage [7] estimated the survival function for a human conceptus throughout gestation by combining information from: studies estimating the probability of fertilization per menstrual cycle in women trying to have children (e.g., [24]), ii) the fate of a sample of 107 ovulated eggs in women who had hysterectomies soon after a potential conception, and hence the fate of the egg could be evaluated by direct inspection [25], iii) combining information from all previous studies of preclinical (occult) abortions detected by elevated levels of hCG (e.g., [15]), and iv) life tables constructed by following the survival of 3,083 clinically detected pregnancies (e.g., [26]). These data were collected predominantly from women in the USA (most of European descent) living under economically developed conditions. When he plotted the combined data, he concluded by inspection that the survival function appeared to follow a curve resembling a “*first-order decay in a mixed population.*” Boklage’s estimate of this survival function was:

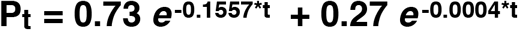

where P_t_ is the probability of a conceptus surviving to time t, and t is days since conception. The value 0.73 is the estimated proportion of high-risk conceptuses (presumed to have abnormal karyotypes) with a rapid decay in survival with age, and 0.27 the proportion of normal-risk conceptuses with much higher daily survival (presumed to have normal karyotypes). Using this equation, the average abortion rate is 0.757. I have denoted this estimate by the circle labeled “B” in Fig. 3, centered over the age of 29 (the average age of women in the study by Wilcox [15]). This value is well above the maximum abortion rate predicted by the data from clinically detected pregnancy rates when sperm is delivered at the optimal time (Fig. 3, curve 5).

An alternative way to make use of the Bokeman equation is to use it to estimate the ratio of occult to clinically detected pregnancies. Assuming clinical pregnancy detection at 21 days post conception, the ratio of occult to clinical pregnancies is 13.15 to one (using a later pregnancy detection time would increase the ratio). Using this ratio, I have estimated the level of occult abortions missing in the Danish study of age-specific abortion and added these to curve 2 (curve 6 in Fig. 3). The curve is close to the estimated ceiling predicted by maximum pregnancy rates at older ages with optimal timing of sperm delivery (curve 5), but consistently above this ceiling at younger ages. However, as noted by Boklage [7], “*Because available data involve such wide methodological variation, and were not in general collected with analysis of the types suggested here in mind, all of the estimates generated here must be considered preliminary.”* The same limitations apply to a similar first-order decay in a mixed population model developed earlier by Wood [5]. Nonetheless, there is strong biological support for the two-category (high and low mortality) model of conceptus mortality because cytogenetic studies of aborted conceptuses consistently find a high proportion that are karyotypically abnormal, and nearly all conceptuses with one or more monosomies or trisomies abort.

To obtain an unbiased and statistically rigorous estimate of the age-specific abortion rate, Darryl Holman and James Wood designed and implemented a study to directly estimate the age-specific abortion rate in a natural fertility population in Bangladesh [1,8-9]. Their analysis allows for the existence of two biologically distinct categories of conceptuses, each with its own distinct survival curve: one type with an abnormal karyotype that causes a rapid, exponential decline in surviving embryos over time (nearly all of these conceptuses would be expected to abort within 6 weeks of age), and another type with a normal karyotype that has a much less steeply declining exponential decay function (due to all other causes of pre-term mortality). To estimate the parameters of these two survival functions, a total of 1,561 menstrual cycles from 708 women were monitored using 4,400 urinary assays for hCG (50% of pregnancies could be detected by 26.3 days since last onset of menses, with the earliest detected pregnancies at about 3 weeks). At least some portion of 329 pregnancies were included in the study. From these data, a maximum likelihood approach was used to simultaneously estimate the false positive and false negative rates of the pregnancy test as well as age-specific: i) fecundability (probability of conception during a potentially fertile menstrual cycle), and the probability of abortion due to ii) a karyotypically normal and iii) a karyotypically abnormal conceptus. Their maximum likelihood estimated curve of age-specific abortion rate is shown by curve 6b in Fig. 3. The predicted abortion rates are consistently and substantially higher than the estimated ceiling predicted from clinical pregnancy rates of European women. This difference feasibly reflects differences between the abortion rates of populations of women experiencing markedly different nutritional, medical, and socio-economic conditions.

To utilize the exceptionally high quality data from the Holman and Wood study, I next focused on their estimate of the proportion of total abortions that were occult vs. clinically detectable. If women in Bangladesh had higher abortions rates because less favorable environmental conditions produced higher abortion rates throughout pregnancy (irrespective of conceptus category –karyotypically normal or abnormal), then the ratio of occult to total abortions would be expected to be far more similar to that of the Danish population compared to total abortion rates. Using these estimates (Fig. 8.17 from Holman 1996), the estimated ratio of total abortions (i.e., clinical + occult) to clinically detected abortions (clinical) was 6.258, 8.318, 9.280, and 9.706 for age 18, 28, 38, and 48 respectively, assuming a clinical detection of pregnancy on day-21 post-conception in the Danish data set. Using a cubic-spline to interpolate intervening values, I have plotted Holman-Wood-adjusted abortion rates for the Danish data set in Fig. 3 (HW-adjusted curve, curve 7, in orange). I consider this curve to be a potentially unbiased estimate of the human age-specific abortion rate in Denmark.

Although the HW-adjusted curve (curve 7 in Fig. 3) falls between the ceiling determined from maximum clinical pregnancy rates given optimal timing of sperm delivery, and the floor determined by adding-back some of the abortions not recorded in the Danish study (i.e., adding abortions that were not associated with hospitalization and some of the occult abortions), a more detailed assessment of the accuracy of the HW-adjusted curve can be achieved with data on the rate of karyotypic abnormalities in early-stage human embryos. Nearly all aborted conceptuses in the HW-adjusted curve of age-specific abortion rate were estimated to be karyotypically abnormal [1,5,8,9]. One can compare the abortion rate predicted by the HW-adjusted curve in Fig. 3 to the rate of karyotypically abnormal early-stage embryos in samples from women of different ages (nearly all karyotypically abnormal embryos will abort, although some conceptuses with trisomy for chromosome 21 [ < 20%, [27]], trisomies for chromosomes 8, 13, and 18 (<< 1%, [28]) and X0 karyotypes [~ 1%, [29]] survive to birth). This type of data comes from three different sources.

The first data source is a one-of-a-kind study in which Labarta et al. [11] collected 46 eggs from egg donors without chemical stimulation (that is routinely used in assisted reproductive technology [ART] clinics to induce multiple ovulated eggs per menstrual cycle). Egg donors are healthy, fertile women who donate eggs to be used by other couples who are experiencing fertility problems. This study therefore eliminates any elevation in chromosomal abnormality rate that may be an artifact of: i) women using ART clinics not being a random sample of the population as a whole, and ii) chemical stimulation of the mother to ovulate multiple follicles during a single menstrual cycle. To guarantee fertilization, each egg was fertilized via intracytoplasmic sperm injection (ICSI), and then a cell from a 3-day old embryo was screened via FISH (fluorescence *in situ* hybridization) for ploidy level for nine chromosomes (13, 16, 18, 21, 22, 15, 17, X, and Y). Sampling a single cell from an embryo may produce an inaccurate estimate of the proportion of karyotypically abnormal embryos due to mosaicism, but in this study the researchers reanalyzed samples of embryos scored as karyotypically abnormal on day-3 (based on a single cell) again on day-5 (scoring every cell in the embryo) and found 92% concordance (i.e, 92% of all day-3 embryos scored as abnormal –by assay of a single cell– were also scored the same two days later when all cells were measured), so this potential problem was not substantial. The proportion of embryos displaying abnormal karyotypes was 34.8% (16 of 46). Munne et al. [30] used a combination of 12 chromosome FISH as well as comparative genomic hybridization (CGH, which measures ploidy at all chromosomes simultaneously with 98.1% accuracy) to show that the 9 chromosome FISH protocol used by Labarta et al. [11] would be expected to detect 72.65% of all karyotypically abnormal embryos: so the true karyotypic abnormality rate in the unstimulated sample is estimated to be 100^*^(.348 / .7265) = 47.9%. The mean age of the oocyte donors (± SD) was 25.4 ± 4.0 years old, so in Fig. 3 I have denoted this estimate for the spontaneous abortion rate (weakly downwardly biased, since it would not include karyotpically normal aborted conceptuses) by a green circle above age 25.4. There is a close match between the estimated proportion of karyotypically abnormal embryos and the HW-adjusted curve for abortion rate (orange curve 7 in Fig. 3).

In a parallel treatment of the same egg donors with a “moderate” dose of gonadotropins for ovarian stimulation (150 IU FSH and 75 IU human menopausal gonadotropin; [11]), the number of eggs analyzed was increased to 303 and the estimated percent of karyotpically abnormal embryos (after adjustment for screening only 9 chromosomes) was 52.6% (which did not differ statistically from the unstimulated sample of 46 eggs, P = 0.64). I have denoted this larger-N estimate of the spontaneous abortion rate (also weakly downwardly biased, since it would not include karyotpically normal aborted conceptuses) by a black circle at age 25.4. Again, there is close agreement with the WH-adjusted abortion rate (Curve 7 in Fig. 3).

The second test of the HW-estimate for age-specific abortion rates comes from a study by Sills et al. [12] on women using in vitro fertilization (IVF). A total of 24 young egg donors (mean age = 24.0 ± 2.7 years, range = 20-29) were used to generate 305 embryos. Of 284 successfully karyotyped embryos (screening all chromosomes simultaneously using array-CGH), aneuploidy was found in 53.2% (Most embryos [86%] were biopsied on day-3 post-fertilization, with the remainder on day-5). I have denoted this estimate for the spontaneous abortion rate (weakly downwardly biased, since it would not include karyotpically normal aborted conceptuses) by a blue circle at age 24. This estimate may also be weakly downwardly biased because array-CGH misses a minority of karyotypic anomalies (see below). Again, there is close agreement with the WH-adjusted abortion rate (Curve 7 in Fig. 3). Potential biases in these data are discussed below.

The third test of the HW-adjusted estimate for age-specific abortion rates is based on a study by Ata et al. [13] which used array-CGH to assay for karyotypic abnormalities in a very large sample (N = 5,918) of cleavage-stage embryos obtained from 726 women across 70 North American assisted reproduction clinics. The rate of abnormal karyotype vs. age from this study is shown in Fig. 3 by the blue dashed curve (curve 8). Again, this estimate may be weakly downwardly biased because array-CGH misses a minority of karyotypic anomalies (see below). Again, there is fairly close agreement with the WH-adjusted abortion rate (Curve 7 in Fig. 3).

The three tests of the HW-adjusted curve for age-specific abortion rate described in the above three paragraphs have potential biases. These biases may be a consequence of: i) intracytoplasmic sperm injection (ICSI), ii) hormonal ovarian stimulation to obtain eggs for assisted reproduction protocols, iii) analysis of women using ART facilities (who may have elevated rates of karyotypically unbalanced eggs), and iv) array-CGH missing some haploid, or triploid of higher ploidy level embryos. A recent large-scale study of the accuracy of array-CGH found that it detected 98.1% of all karyotypic anomalies [31], so any bias from the use of array CGH should be minor. There is empirical evidence that ovarian stimulation (used to generate eggs for in vitro fertilization) increases aneuploidy rate (e.g., [32]), although other studies found no effect (see summary in [33]), so the estimates for embryonic aneuploidy used to check the accuracy of the WH-adjusted abortion curve (curve-7 in Fig. 3) may be biased upward. However, the collective effect of this factor and the other potential biases are expected to be small because a recent meta-analysis of the rates of chromosomal anomalies from 15 studies with a collective sample size of 1,896 aborted conceptuses from ART mothers and 1,186 controls (no ART) mothers, found no difference in abnormal karyotype rate as well as no difference in this rate between IVF and ICSI categories, nor between hormonally stimulated and no stimulation categories [34]. This meta analysis indicates that any biases in the studies that I have used to verify the spontaneous abortion rate predicted by the HW-adjusted curve are small. This conclusion is reinforced by the high similarity of four independently estimated rates of karyotypically abnormal embryos for women in their mid-twenties with different combinations of level of hormonal stimulation and donor vs. ART-mother origin of eggs (curve-8, blue, green and black points in Fig. 3).

The concordance between the estimated abortion rate depicted in curve-7 (orange) of Fig. 3 (i.e., the HW-adjusted estimate of the age-specific estimated abortion rate) and the three independent estimates of age-specific rates of abnormal karyotypes in early-stage embryos provides strong evidence that the HW-adjusted estimate of the abortion rate is a close approximation to the true abortion rate in an economically developed population. Hereafter I will refer to this estimated curve as the HW_adj_- curve.

### Age-specific abortions per newborn

The age-specific average number of abortions that occur during the production of a live birth (i.e., the abortion cost per human newborn) can be estimated from the HW_adj_-curve. During each menstrual cycle, one follicle is drawn via hormonal stimulation (i.e., via natural secretion of FSH) from a very large pool of dormant follicles. After fertilization, the outcome of each conceptus is a Bernoulli trial (success or failure) in which the height of the HW_adj_-curve is the estimated probability of an abortion, and its complement is the estimated probability of a live birth. It is reasonable to assume that –in women with normal fertility– the fate of each conceptus is independent of the fate of previous conceptions because its fate depends predominantly on its karyotype –which is determined by a new draw from the pool of dormant oocytes and and a new draw from the sperm donor. In this case, the number of failures (abortions) that occur before the first success (newborn) will follow a negative binomial distribution (NB[r,p]) with parameters r = 1 = one live birth, and p = the probability of a newborn, i.e. number abortions per newborn ~ NB(1,prob_newborn_). The mean of a NB(1,p) distribution is its odds ratio = (1-p)/p and its variance = (1-p)/p^2^. For example, at age 32, the HW_adj_-curve predicts a 60% probability of a conception ending in an abortion, so the average number of abortions per newborn is 0.6/0.4 = 1.5 and the variance 0.6/0.4^2^ = 3.75 and the standard deviation = √3.75 = 1.94. Using this relationship, I have plotted the expected (i.e., the average) number of aborted conceptions per newborn across all maternal ages (Fig. 4).

**Figure 4.**
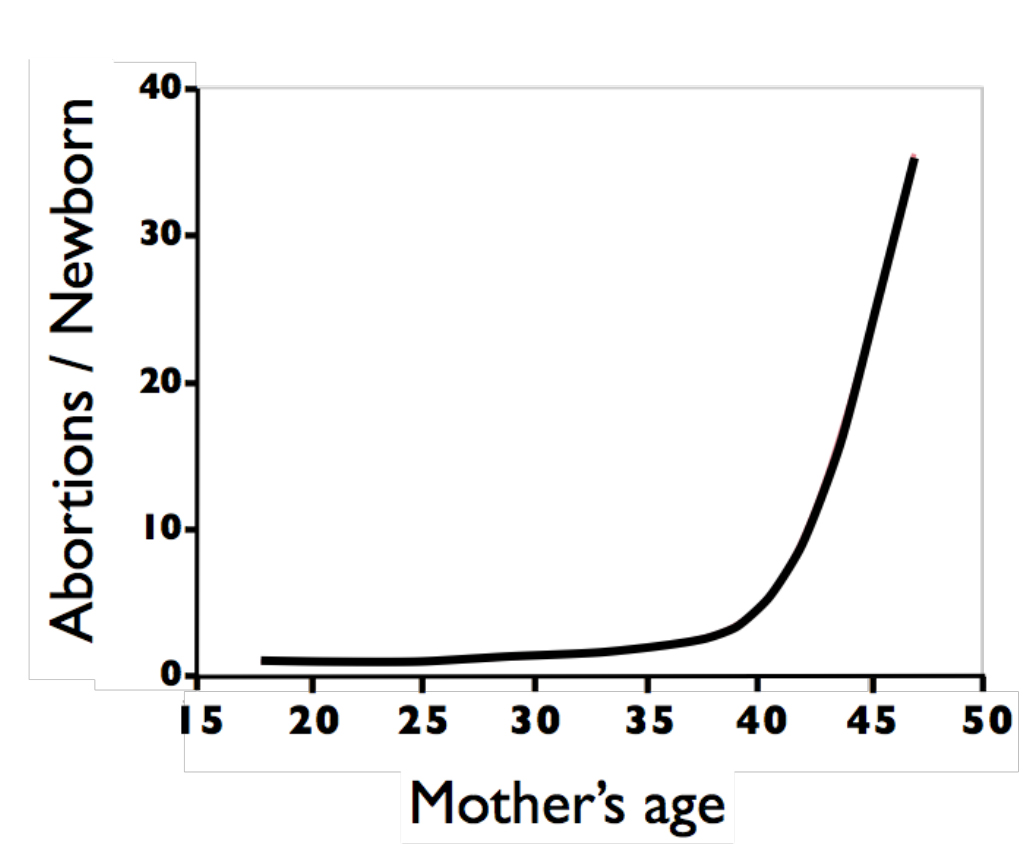
The estimated age-specific ratio of the average number of spontaneous abortions that are produced during the production of a newborn.

### Abortions per newborn in frontier Mormons in the mid 19^th^ century and Danes at the end of the 20^th^ century

Fig. 4 can be used to determine the average number of abortions per lifetime. An exact calculation would require the probability distributions for a prohibitively large number of age-specific parameters, e.g. marriage, frequency and timing of sexual intercourse, birth intervals, duration of lactational amenorrhea, and so on. To my knowledge, such a complete set of accurate information is not available for any human population. However, for the purpose of comparison, abortions per lifetime can be calculated for a “canonical” woman whose marriage age, birth intervals, and lifetime number of children are all constrained to be the population norms (i.e., average values). The logical basis for constructing a canonical woman is the same as that for constructing a canonical molecular sequence for a polymer –such as DNA, RNA, and protein– that has the most prevalent unit at each position in a contiguous sequence: it provided a standardized reference point for comparisons. Few women will be truly average for all these parameters simultaneously (as is true for polymers and their canonical sequence), but this calculation nonetheless provides a basis for comparison.

One of the most extensive data sets for women following a natural fertility mode comes from the detailed records kept by frontier Mormons during their population expansion in the Western United States. From these records the distribution of family sizes and inter-birth intervals for these families has been tabulated by Anderton & Bean [35]. Using the average marriage age of a woman, average birth intervals and family size during a period when the population remained under natural fertility conditions (the 1840s) reported in [36] and [35], the abortions per lifetime can be calculated for a “canonical” frontier Mormon woman. This value is the sum of eight NB(1,pi) values, where pi is one minus the expected abortion rate taken from the HW_adj_-curve in Fig. 3 (curve 7). This sum is 16.8 abortions per lifetime. It is reasonable to apply the HW_adj_-curve from Fig. 3 to frontier Mormons because this population was composed primarily of people with western European ancestry (including many Danes and other Scandinavians) and because this population was well nourished [37].

If I had included variation in marriage age and birth intervals, the average of 16.8 abortions per lifetime would be increased because the curve of abortions per newborn has a positive second derivative, causing random variation to earlier times to have a smaller reducing effect on lifetime abortions than the increasing effect of random variation to later times. This is especially relevant for the last few children produced because of the steep rise in abortions per newborn after age 35 (Fig. 4). By constraining the “canonical” woman to have the average birth interval between each of eight newborns, a woman’s last conception occurs at age 39. If a last child were produced during the early forties, or sexual intercourse continued into this time period after a last child was produced in her late thirties, the high ratio of abortions per newborn at these elevated ages would cause a substantial increase in her abortions per lifetime. So, the value 16.8 abortions/lifetime reported here for a “canonical” frontier Mormon woman is conservative –probably to a high degree.

The negative binomial distribution has a high variation when the probability of success (p) is small (variance = [1-p]/ p^2^]), and this leads to substantial variation about the mean value of abortions per lifetime reported in the above paragraph. To estimate the variation about the mean value of 16.8 abortions per lifetime for a canonical woman, I have generated 100,000 random samples of eight NB(1,p_i_) values (with p_i_ determined from Fig. 3 using the average ages at marriage and birth of each of 8 children (taken from [36] and [35]) and then tallied their sum in Fig. 4, red histogram. This distribution of abortions per lifetime for a canonical frontier Mormon woman (constrained to have average age at marriage, family size and birth intervals) shows that it would not be uncommon (i.e., >20% of the time) for a woman following a natural fertility birth pattern to have more than 23 abortions per lifetime.

For comparison, I calculated the abortions per lifetime for a canonical Danish woman –who would have had free access to modern birth control and elective abortions up through week 12 of gestation– during the time frame when the abortion data reported in Figs. 1 and 3 were measured (Fig. 5, blue histogram). These woman had an average family size of 1.7 children with an average of 0.37 elective abortions per newborn [38-41]. The distribution in Fig. 5-blue (including 0.37 elective abortions per newborn added as a binomial random variable) has a mean of 2.51 abortions per life time, and the 95% upper bound of the distribution (6.3 abortions per lifetime) is far less than the average for the frontier Mormon women, i.e. the frontier Mormons had 570% more lifetime abortions. So modern birth control, even when elective abortions of early-stage conceptuses are legally and freely available, leads to markedly fewer abortions per lifetime compared to natural fertility.

**Figure 5.**
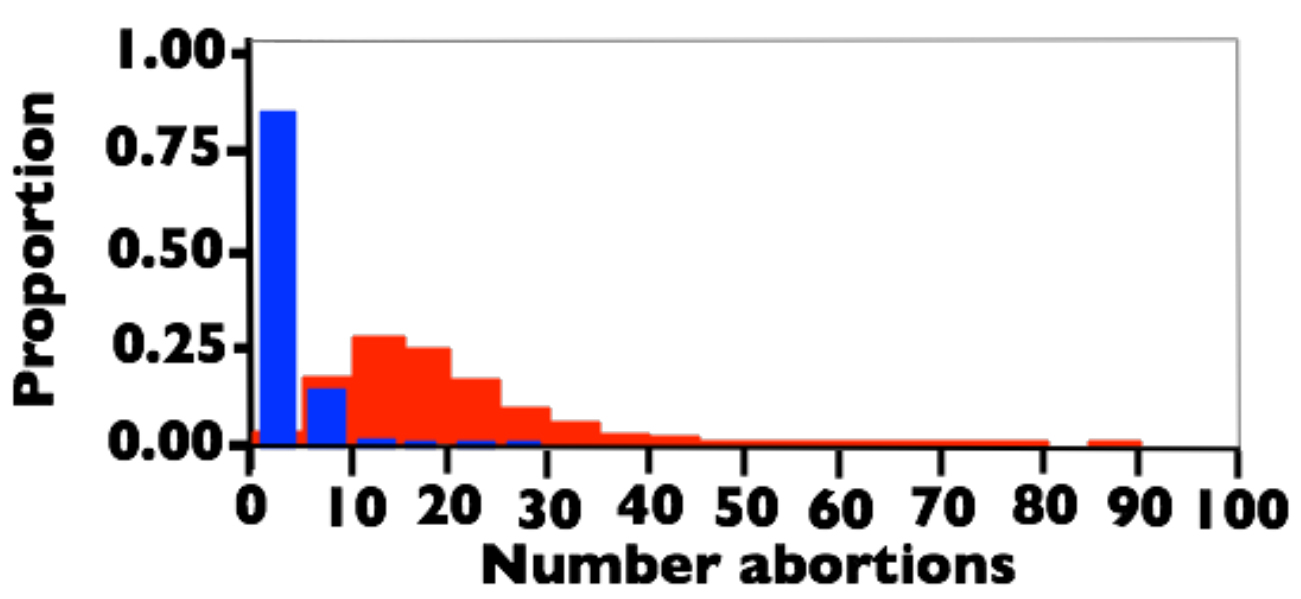
The estimated distribution of abortions per lifetime for a canonical woman constrained to have average age at marriage, age at first birth, birth intervals, and family size. Red: Frontier Mormons during the 1840s. Blue: Danes between 1978-1992. See text for sources used to estimate the demographic parameters.

One of the striking features of the calculations displayed in Figs. 3-5, and a conclusion reached in earlier studies [1,5,7-9,15] is that the predominant outcome of fertilization in a human female adhering to a natural fertility program is abortion: in my canonical analysis of frontier Mormons, on average, abortions outnumbered newborns by more than a factor of two (16.8 to 8).

## Discussion

Abortion, in the form of a miscarriage during a clinically detected pregnancy, is the exception rather than the norm, when one excludes older women or those with a predisposing medical condition [4]. This observation leads to the intuitive conclusion that spontaneous abortion is a fairly uncommon reproductive anomaly in humans. However, studies in the 1980s using highly sensitive assays for cGH (e.g. [15]) clearly demonstrated that most spontaneous abortions are occult and go completely unnoticed by women. Subsequent studies estimating total abortion rate [1,5,7-9] made a strong case that spontaneous abortions are common in humans, even at young ages. The most comprehensive data, including maximum likelihood estimates of all relevant parameters, came from a natural fertility population in Bangladesh [1,8,9]. The data that I surveyed in this study expands the previous work by integrating many very large data sets to estimate the age-specific abortion rate (and test the estimate’s accuracy) in a near-contemporary population from an economically developed country (the HW_adj_-curve; orange curve-7 in Fig. 3). This analysis corroborates the previous studies and thereby empirically reaffirms the conclusion that spontaneous abortion is a common and fundamental component of human reproduction at all ages.

The estimated curve of abortion rate vs. age reported here (HW_adj_-curve) differs from the previous point estimate based primarily on data from USA women ([7]; the point labelled “B” in Fig. 3) and the age-specific curve from a natural fertility population in Bangladesh [1,8,9]. The difference from the USA estimate is not surprising because Boklage [7] averaged across all ages and because he used data collected for other purposes – so his estimate was therefore only meant to represent a first approximation. The Bangladesh estimate does not share the problems found in the USA estimate, so the difference between this estimated curve (curve 6b in Fig. 3) and the one reported here (HW_adj_-curve, curve-7 in Fig. 3) is probably real. However, there is no *a priori* reason to expect spontaneous abortion rates to be identical between populations experiencing markedly different nutrition, socio-economic conditions, and access to modern medicine –and as would be expected, the spontaneous abortion rate is estimated to be higher in the population experiencing less optimal conditions.

To take advantage of the rigorous statistical analysis used to estimate the age-specific abortion rate in the Bangladesh study, I focused on the estimated proportion of total abortions that were occult. There is no guarantee that this value will be the same, or similar, in the Bangladesh and economically developed populations, but it was a plausible outcome. Making this assumption, I generated my estimate for the age-specific abortion rate among Danes by adjusting the Andersen et al. data [4] for clinically observed abortions) using the ratio of occult to clinical abortions observed in the Bangladesh data set [1]. The accuracy of this curve was supported by several independent cytological studies screening many thousands of embryos (a collective sample size of over 6.5K), that estimated rates of karyotypic abnormalities in early-stage embryos. The close correspondence between the cytological data and the estimated age-specific abortion rate makes a strong case that this estimate (curve-7 in Fig. 3) is a close approximation to the true rate.

The estimated curve for age-specific abortion rate can be used to estimate the cost, in the currency of spontaneous abortions, of producing a newborn (Fig. 4). The curve shows why, in humans, a choice to produce one or more newborns must come at a considerable abortion cost. Abortion is nearly as common as live-birth for conceptions that occur in a woman’s early-twenties, but after the mid-twenties, abortions are the norm rather than the exception.

## Conclusions

A synthesis of many large-scale studies from the last 15 years unambiguously confirms the Wood-Boklage-Holman hypothesis that abortion is an intrinsic and overarching component of human reproduction. It is the most common outcome of conception across a woman’s lifetime and the predominant factor controlling age-specific variation in human female fertility. To reproduce, a human female cannot forgo a high risk of abortion, and to have a large family it is virtually impossible to avoid multiple abortions. Modern birth control with access to elective abortions, markedly reduces –rather than increases– the lifetime number of abortions a woman produces.

